# Deciphering hybrid larch reaction norms using random regression

**DOI:** 10.1101/301119

**Authors:** Alexandre Marchal, Carl D. Schlichting, Rémy Gobin, Philippe Balandier, Frédéric Millier, Facundo Muñoz, Luc E. Pâques, Leopoldo Sánchez

## Abstract

The link between phenotypic plasticity and heterosis is a broad fundamental question, with stakes in breeding. We report a case-study evaluating temporal series of wood ring traits of hybrid larch (*Larix decidua* × *L. kaempferi* and reciprocal) in relation to soil water availability. Growth rings record the tree plastic responses to past environmental conditions, and we used random regressions to estimate the reaction norms of ring width and wood density with respect to water availability. We investigated the role of phenotypic plasticity on the construction of hybrid larch heterosis and on the expression of its quantitative genetic parameters. The data came from an intra-/interspecific diallel mating design between both parental species. Progenies were grown in two environmentally contrasted sites, in France. Ring width plasticity with respect to water availability was confirmed, as all three taxa produced narrower rings under the lowest water availability. Hybrid larch appeared to be the most plastic taxon as its superiority over its parental species increased with increasing water availability. Despite the low heritabilities of the investigated traits, we found that the quantitative genetic parameters varied along the water availability gradient. Finally, by means of a complementary simulation, we demonstrated that random regression can be applied to model the reaction norms of non-repeated records of phenotypic plasticity bound by a family structure. Random regression is a powerful tool for the modeling of reaction norms in various contexts, especially perennial species.

## INTRODUCTION

Heterosis in plants is a phenomenon that usually arises in unfavorable environments, as observed in a wide range of crop hybrids, like corn (Janick 1999; Gallais 2009). The relative stability of hybrids when the parents’ performances drop is called ‘hybrid homeostasis’ and, as a consequence, the hybrids are often expected to perform better than their parents in most environmental conditions. The stability of heterosis across sites has recently been highlighted for hybrid larch (*Larix decidua* Mill. × *L. kaempferi* (Lamb.) Carr., and the reciprocal cross), a highly productive conifer cultivated for wood in Western Europe and North America (Marchal *et al.* 2017). For this reason, phenotypic plasticity is suspected to play a key role in the construction of hybrid larch (HL) heterosis.

Phenotypic plasticity, in its narrow-sense definition, is the ability of a genotype to produce several phenotypes depending on the environmental conditions. It can be studied by means of reaction norms (Schlichting and Pigliucci 1998). A reaction norm is an equation, or simply a graphical representation, of the value taken by a phenotype along an environmental gradient. Estimating reaction norms is not a trivial task. In particular, exposing the same genotype to different environments may be experimentally challenging, depending on the biological model. In some cases, the only solution is to expose related individuals to the different environments, relying on their genetic relationship to draw the common, additively inherited, component of the reaction norms (*e.g.* Gibert *et al.* 2004; Valladares *et al.* 2006). The random regression model is an extension to the classical quantitative genetics model (Kirkpatrick and Heckman 1989), and as such it can pre diet the additive component of reaction norms, and give access to causal components of population variation. In addition, Murren *et al.* (2014) demonstrated with a large meta-analysis that when comparing reaction norms of close species or populations, changes in shapes (*i.e.* slope, curvature) were generally higher than the changes in the intercepts (*i.e.* the taxon means), suggesting the need of high-order modeling. In that view, random regression modeling has also the valuable capability to fit complex curves for reaction norms.

Random regression is a special case of covariance function (Meyer 1998). Covariance functions present a particular interest in quantitative genetics, as they allow the representation of quantitative genetics parameters as functions. For instance, the cattle breeding literature is rich in illustrations of heritabilities and genetic correlations estimated as functions of time in the milk production context (*e.g.* Miglior *et al.* 2007; Muir *et al.* 2007; Jamrozik *et al.* 2010). Although covariance functions and random regression have been often suggested for the modeling of reaction norms (Kirkpatrick and Heckman 1989; De Jong and Bijma 2002; Schaeffer 2004), their application is still rare for most taxa (Morrissey and Liefting 2016). For instance, in the forestry context, random regression has been used to model growth over time (Apiolaza and Garrick 2001; Wang *et al.* 2009), but not tree growth reaction norms over environmental gradients until very recently (Carnwath and Nelson 2016; Marcatti *et al.* 2017). Li *et al.* (2017) advocated for the use of random regression in forest trees, but the growth reaction norms they reviewed were only performed at the population scale, with fixed effects, without covariance functions. Nevertheless, because they are sessile, long-living, and because they record radial growth increments in the form of annual rings, trees are remarkable biological models for the study of phenotypic plasticity. The use of random regression has been suggested for the study of tree rings plasticity (Sánchez *et al.* 2013).

For a given tree, the succession of wood annual rings constitutes an archive of its plastic responses to the succession of past climatic environments. Indeed, trees growth is conditioned by solar radiation, temperature and precipitation. They are particularly sensitive to drought, as this can eventually lead to lethal embolism of their hydraulic system. As a response to water stress, trees have a range of plastic responses including modification of the cambial activity to control the sap flow rate and xylem resistance to embolism (Bréda *et al.* 2006; Rennenberg *et al.* 2006). Under a temperate climate, water availability is usually high in spring with cool temperatures and lower in summer with higher temperatures. Trees respond to this seasonal succession by producing an early wood, made of large cells with thin walls, progressively followed by a late wood, made of narrow cells with thick walls, allowing a rapid change in the trunk of the water conductance along the season (Sánchez-Vargas *et al.* 2007; Martinez-Meier *et al.* 2009; Bryukhanova and Fonti 2013). This succession of early and late wood is recorded in the wood as an annual growth ring.

In summary, trees present the remarkable feature of recording long repeated series of phenotypes that are known to respond plastically to the climatic environment, and differentially among individuals. This provides a gold mine of information for the study of phenotypic plasticity. First, based on a random regression approach, we leveraged this information to address the question of the role of phenotypic plasticity in the construction of HL heterosis for radial growth. Indeed, integrative heterosis as observed for HL stem circumference necessarily arises from the cumulation of heterosis occurring at the annual ring scale: this might be expressed differentially with respect to parental genotypes depending on water availability. Secondly, we described how the water availability affected the quantitative genetic parameters of HL and of its parental species. Finally, since most biological models do not archive repeated series of phenotypic plasticity records, we addressed the question of the generalizability of the method. Using simulations, we evaluated the robustness of the random regression approach for the modeling of reaction norms in the case where no repeated series of phenotypes per individual are available.

## MATERIALS AND METHODS

The data are part of a multi-site progeny trial established in early 1997 on two environmentally contrasted sites at Saint-Appolinaire (SA, 45°58′N 4°26′E, 784 m a.s.l.) and Saint-Saud (SS, 45°31‱N 0°48′E, 307 m a.s.l.) in central France. The site of SA is a relatively high-elevation site, on a steep slope with a southern aspect formerly planted with Douglas-fir; whereas the site of SS is a low elevation site on a former meadow, under oceanic influences. Progenies were produced by control-crossing in the frame of a diallel mating design between 9 European larch (*Larix decidua*, EL) parents and 9 Japanese larch (*L. kaempferi*, JL) parents, producing pure species and HL full-sib progenies. The set-up is described further in Marchal *et al.* (2017).

### Phenotypic data: wood formation records

One breast-height diameter increment core was collected from each tree from each site. For each diameter increment core, only one radius (the one exhibiting the fewest defects) was kept for further analysis. These radial increment cores were sawed in 2 mm thick boards and X-rayed to obtain microdensitometric profiles (Fig. 1). From the alternating of early wood and late wood, the year of formation for each ring was identified (Regent Instruments Canada Inc. 2008). Ring width (RW) and ring mean density (RMD) were measured from the microdensitometric profiles. A total of 1998 increment cores in SA and 2278 increment cores in SS were collected. In SS, the increment cores were collected at three different periods: before the first thinning in 2003 from trees to be felled, and two later ones before and after the second thinning, that is in 2009 and in 2011. In SA, the collection was done in 2013. The average number of rings available per core was 13.2 in SA and 9.7 in SS (Supplementary 1, Fig. S 1).

**Figure 1.**
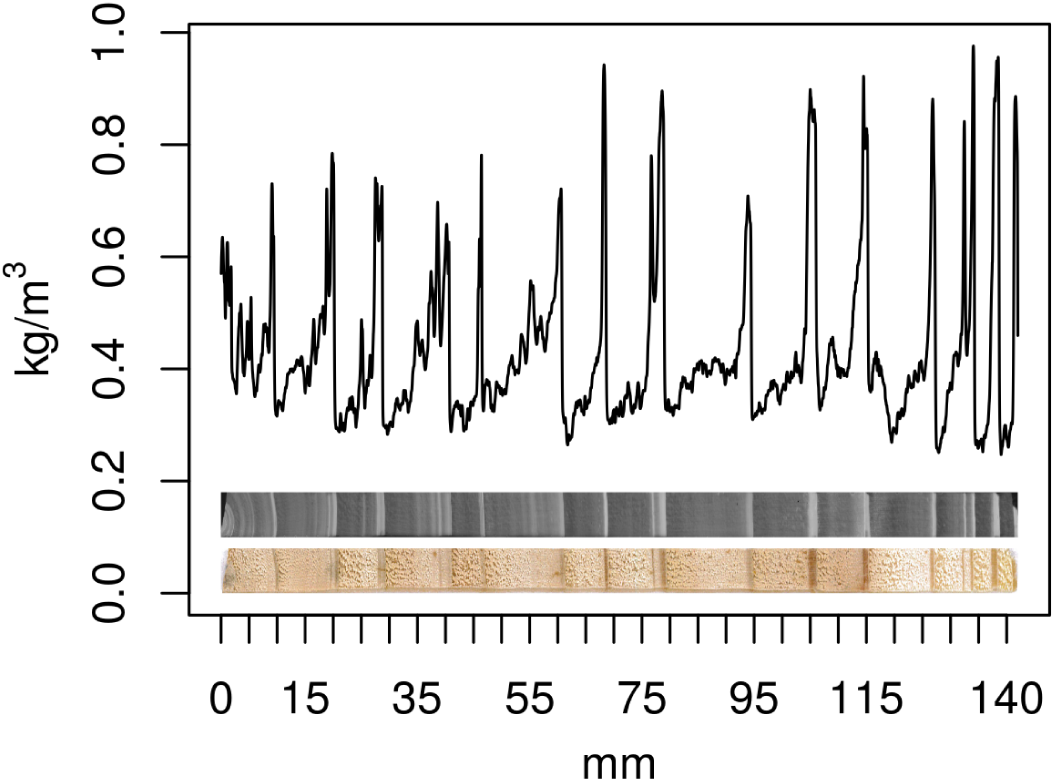
Microdensitometric profile from one wood core sample in Saint-Appolinaire. Bottom: wood core sample (pith on the left). Middle: X-ray radiography of the sample. Top: wood density variation (kg/m^3^) along the core (mm). The peaks correspond to late wood and delimit the annual growth rings

### Environmental data: soil water availability

The soil daily relative extractable water (REW) content was estimated using a water balance model. The REW varies between 1 (field capacity) and 0 (permanent wilting point, meaning the water is no longer available for the plants), and it was estimated for each combination of site and year for which ring data were available. We used an adapted, simplified implementation of Granier et al. (1999)’s daily water balance model. This implementation allowed us to account for temporal evolution of the stand, and to run the model at the individual scale. Indeed, during the growth of the trees, the canopy leaf area increases and subsequently increases the transpiration, whereas understorey shrubs and grass vegetation decrease. Also, the intra-site spatial heterogeneity of the soil storage capacity causes variation in the water availability at the individual tree scale. The water balance model is described in Supplementary 2. The output of this model is the daily REW available at the tree level.

To summarize a year of fluctuation of REW, we selected the lowest decile of the whole distribution of the year’s daily REW (D1rew). This index was considered because of its easy interpretation (the REW below which lay the ∼35 driest days of the year), because of its likelihood to carry the information of drought events and therefore to affect the growth, and finally because it ensures a proper coverage of the environmental gradient (Supplementary 1, Fig. S 2), unlike e.g. indexes based on a drought threshold that may not be reached some humid years. The environmental gradient coverage guarantees the stability of the regression parameters and the quality of the subsequent analyses.

**Figure 2.**
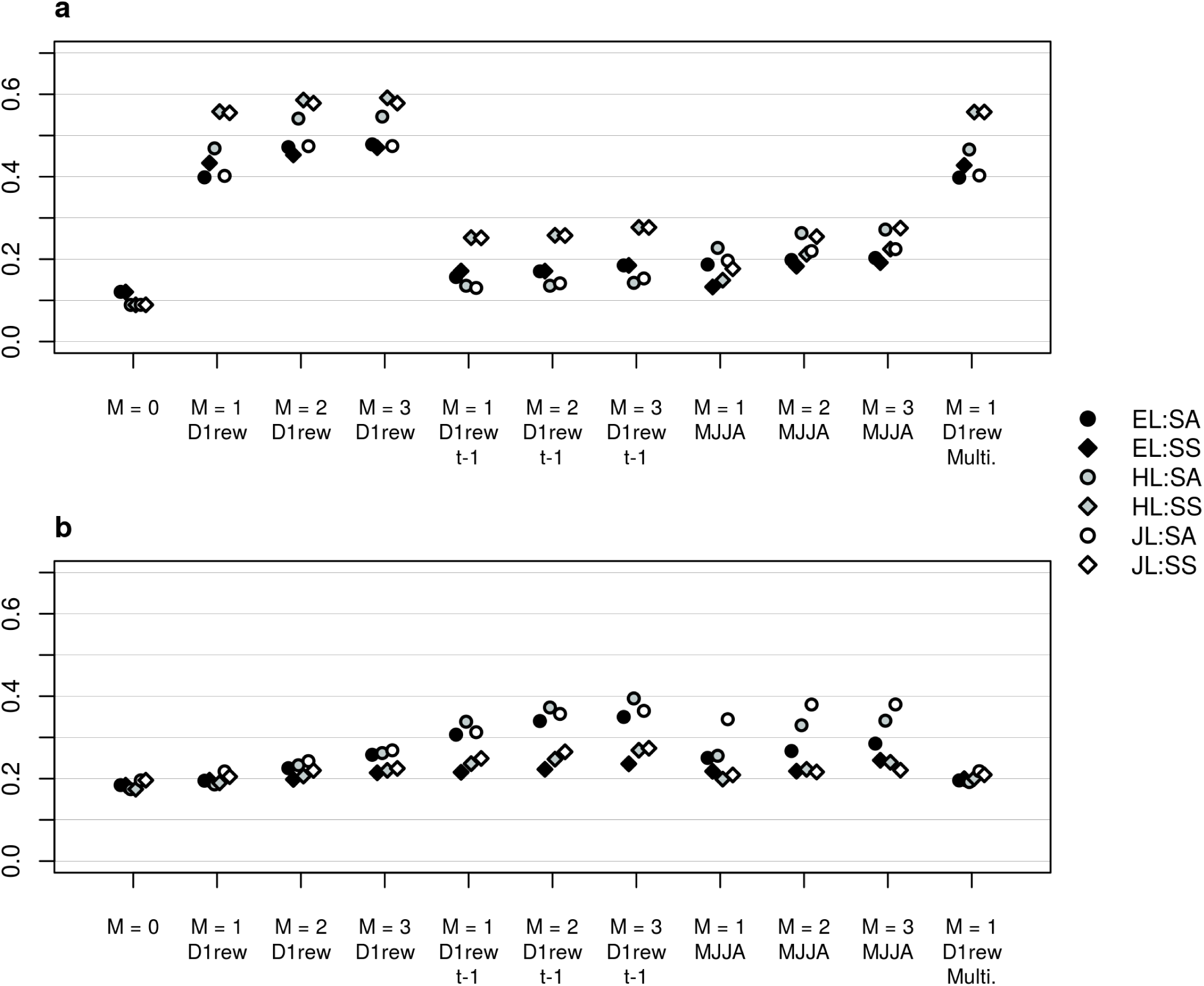
Coefficients of determination (*R*^2^) for ring width (RW) (a) and ring mean density (RMD) (b), depending on the model specification. The polynomial order (*M*) varied from 0 to 3. Three covariates were compared: the first decile of the soil daily relative extractable water (D1rew) for the current year (year *t*, implicit) and for the previous year (year *t* − 1, specified), and the sum of daily differences between precipitation and potential evapotranspiration from May to July (MJJA) for the current year. The last model specification (‘Multi.’) corresponds to 1^st^ order regression with the covariate D1rew of the current year *t*, but in this case the model was multivariate and RW and RMD were analyzed simultaneously

The index D1rew, derived from a complex water balance model, was compared to a simpler index MJJA defined as the sum of rain from May to August (inspired from Fallour-Rubio et al. (2009)), from which was subtracted the daily potential evapotranspiration (Supplementary 1, Fig. S 2). Moreover, Bryukhanova and Fonti (2013) showed that for European larch, several traits including RW were more strongly correlated to the soil water deficit of year *t* − 1 than to the deficit of the current year *t*. Therefore, we used D1rew for year *t*, for year *t* − 1, and MJJA for year *t* as three separate environmental gradients, and we modeled the reaction norms along these environmental gradients separately.

### Modeling of reaction norms by random regression model

Reaction norms were modeled using orthogonal Legendre polynomials (Kirkpatrick *et al.* 1990; Schaeffer 2004). Let *L_m_* (*x′*) be the *m*^th^ order Legendre polynomial of *x′*, with *x′* the standardization of *x* on [−1,1], that is *x′* = 2 * [*x* − *min*(*x*)]/[*max*(*x*) − *min*(*x*)] − 1. We fitted the following model for each taxon (EL, HL or JL):

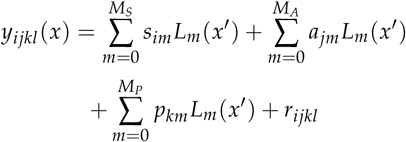

where *y_ijkl_* (*x*) was the *l*^th^ observation of individual *k*, of genotype *j*, from site *i*, with environment *x*. The site’s effect *s* was fixed. The additive effects *a* were random, and depended on the species. For EL and JL pure species, respectively, *a_E_* ∼ *N*(0, **∑_AE_** ⊗ **A_E_**) and *a_J_* ∼ *N*(0, **∑_AJ_** ⊗ **A_J_**); and for the hybrid *a_H_* = *g_E_* + *g_J_* with *g_E_* ∼ *N*(0, 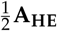) on the EL side and *g_J_* ∼ *N*(0, **∑_HJ_** ⊗ 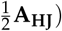 on the JL side (Stuber and Cockerham 1966). Given that the parents were supposed outbred and unrelated, **A_HE_** and **A_HE_** reduced to identity matrices. The permanent environment was *p* ∼ *N*(0, **∑_p_** ⊗ **I_p_**). The residual was unstructured 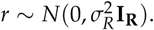 We chose to fix *M_S_* = *M_A_* = *M_p_* = *M* so that an ‘order *M*’ applied to the whole model. Models of different orders are characterized by how much information on plastic responses is included: order 0 does not estimate any plasticity, order 1 calculates slopes, order 2 additionally fits parabolas, and so on.

This model was fitted for both traits RW and RMD, and also for RW and RMD simultaneously in a multivariate approach. In this latter case, the effects *s*, *a* and *p* were estimated for each trait; the variance-covariance matrices **∑** gathered the variances of each combination of trait and order and the covariances between them all; the residual variance 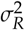 was independent for each trait.

The model was fitted by Markov chain Monte Carlo (MCMC) with the same priors as in Marchal et al. (2017) (Hadfield 2010; R Core Team 2017): parameter expansion was used on the genetic variance-covariance matrices, and flat improper priors were set on the permanent environment variance-covariance matrices as well as on the residual variance. All chains were 5.5 × 10^6^ iterations long, with 5 × 10^5^ iterations burn-in and a thinning of 5 × 10^3^. Point estimations from chains were maximum *a posteriori*, and 95% credible intervals (CIs) were computed where appropriate. The quality of fitting was assessed using coefficients of determination *R*^2^ (Nakagawa and Schielzeth 2013; Johnson 2014).

### Estimation of genetic parameters

The additive variances 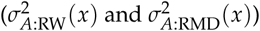 and covariance between RW and RMD (*C_A_*(*x*)) were calculated from the multivariate random regression model as functions of the environmental gradient *x*. The additive variance-covariance matrix decomposes into 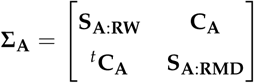 where **S_A:RW_** and **S_A:RMD_** are the sub-matrices for each trait and **C_A_** is the across-traits covariance sub-matrix. The additive variance of a trait *T* is the variance of the additive performances within the population for this trait, that is the variance of a linear combination:

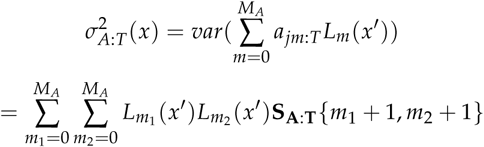

The permanent environment variance
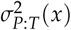
was computed in the same way. Similarly, the additive covariance between the 2 traits was computed as:

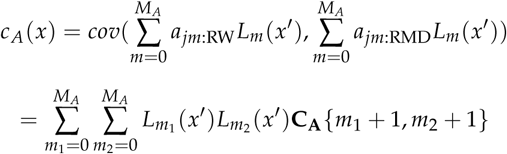

Finally, the permanent environment covariance *c_p_* (*x*) was computed in a similar way. Narrow-sense heritabilities and additive correlation were then computed respectively:

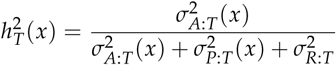

and:

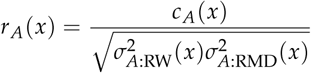

For hybrids, narrow-sense heritabilities and additive correlation were computed for each of the parental contributions *g_E_* and *g_J_*. Therefore, on the EL side:

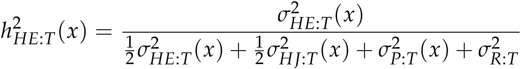

and:

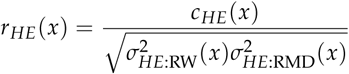

and idem on the JL side.

### Simulation: evaluation of the random regression model with single record per individual

We used the software Metagene to simulate data. The functioning of this simulator to derive *in silica* populations with phenotypic plasticity records is detailed in Supplementary 3. Basically, the genotypic effect at a given locus was set as a function of the environment *α*(*x*) = *α*_0_ + *α*_1_ (*x* + *δ*) + *α*_2_(*x* + *δ*)^2^, where parameters *α*_0_, *α*_1_, *α*_2_ and *δ* defined the parabola that was associated to each genotype in a set of *X* diallelic loci constituting the genome. Then, for each independent simulation, the simulator randomly sampled a genome for each founder, produced the mating between founders, the new offspring genomes, and returned their phenotypic plasticity in the form of longitudinal records over *ϒ* environments. We parameterized the allelic effects in such a way that the additive reaction norms were very interactive, that is, the ranking of the parents varied along the environmental gradient due to slopes and parabolas.

The mating design consisted of a full diallel between 10 monoecious founders, excluding selfs. Each combination of parents (A×B) produced *n*/2 sibs, so that the size of a full-sib family (A×B + B×A) was *n*. The environmental gradient was divided in *n* random positions, each position being defined by an environmental value *x*. Each of these environments hosted one sib per family, and as many individuals as families. The progenies’ phenotypes, their pedigree, and the environmental values *x* were included in the analysis.

We tested 4 different scenarios, resulting from the combination of low (0.1) and high (0.6) heritabilities with small (20) and large (120) family sizes. Thus, these scenarios measured the importance of the quantity (*i.e.* the number of progenies) and quality (*i.e.* the heritability) of information for genetic inference. For each scenario, 100 independent simulations were run and analyzed.

We used the pure species univariate random regression model previously described, without the permanent environment component (as no permanent environment was simulated) and without the site effect (thus the fixed part of the model was a common linear combination of Legendre polynomials of order *M*). The chains were 1.5 × 10^5^ iterations long, with 5 × 10^4^ iterations burn-in and a thinning of 10^3^. We measured the ability of the random regression to infer additive components of the parental reaction norms. To do so, we defined the accuracy of the model as the correlation between the parents’ predicted additive performances and their true simulated additive performances at each point of the environmental gradient. The reaction norms of the parents were only inferred for the ranges on which the progenies were tested.

### Data availability

Our datasets are being submitted to the INRA repository GnpIS (https://urgi.versailles.inra.fr/Tools/GnpIS). Until then, the data are available upon request.

The Metagene simulator is available on the NOVELTREE project page (http://www.igv.fi.cnr.it/noveltree/) and the code is being submitted to GitHub (https://github.com).

## RESULTS

### Quality of model fitting to real data

The order 0 model (fixed and random intercepts) explained between 8.9% and 12.1% of RW variance depending on taxa and site combination (Fig. 2). The addition of fixed and random slopes (1st order) along D1rew of the current year (*t*) greatly improved the model, allowing it to account for 39.9% (EL in SA) to 55.8% (HL in SS) of the variance. Analyzing higher orders (order 2 and order 3 along D1rew of the current year) slightly increased the *R*^2^ (by 5.1% on average compared to order 1). Such high *R*^2^ were not reached using D1rew of the previous year, nor using the simpler index MJJA. For RMD, on the contrary, order 0 or higher orders led to very close *R*^2^ (Fig. 2). Using D1rew of the previous year instead of that of the current year led to a marginal improvement of *R*^2^ that happened only in SA, reaching up to 39.4% for HL with order 3.

Heterosis expressed mostly in radial growth, that was better explained by D1rew of the current year. For this reason, we focused on this environmental gradient only. The gain in *R*^2^ with increasing orders was overall due to more variance explained by the fixed component of the models, as shown in Fig. 3. Therefore, we present in Fig. 4 order 3 reaction norms at the taxon scale (*i.e*. the fixed component of the model), but for parsimony consideration, the multi-trait random regression model (RW and RMD simultaneously), from which the genetic variance parameters were estimated, was fit with order 1. The *R*^2^s of the multi-trait model for each trait were also presented in Fig. 2 and were very similar to their univariate counterparts.

**Figure 3.**
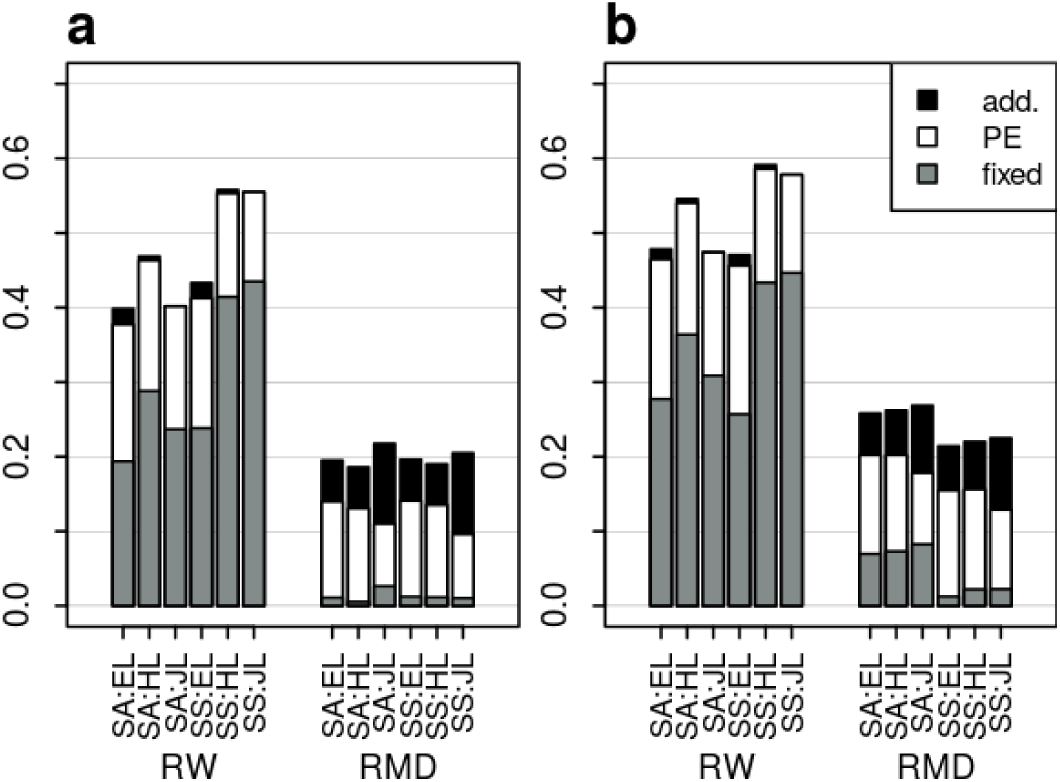
Decomposition of the coefficients of determination (*R*^2^) obtained with the order 1 model (a) and with the order 3 model (b), for each trait: ring width (RW) and ring mean density (RMD), and for each combination of site (SA or SS) and taxon (EL, HL or JL). The environmental gradient was the first decile of the daily relative extractable water for the current year. The proportion of variance explained by each component is presented: genetic additive effects (in black), permanent environment (in white), and fixed terms (in grey)

### Stability of the heterosis and its dependence on the water availability

The ranking in RW performance between EL and JL varied depending on the site: EL performed better in SA while JL did better in SS (Fig. 4). However, the superiority of the hybrid over both its parental references occurred in the two sites, and over the whole range of D1rew.

**Figure 4.**
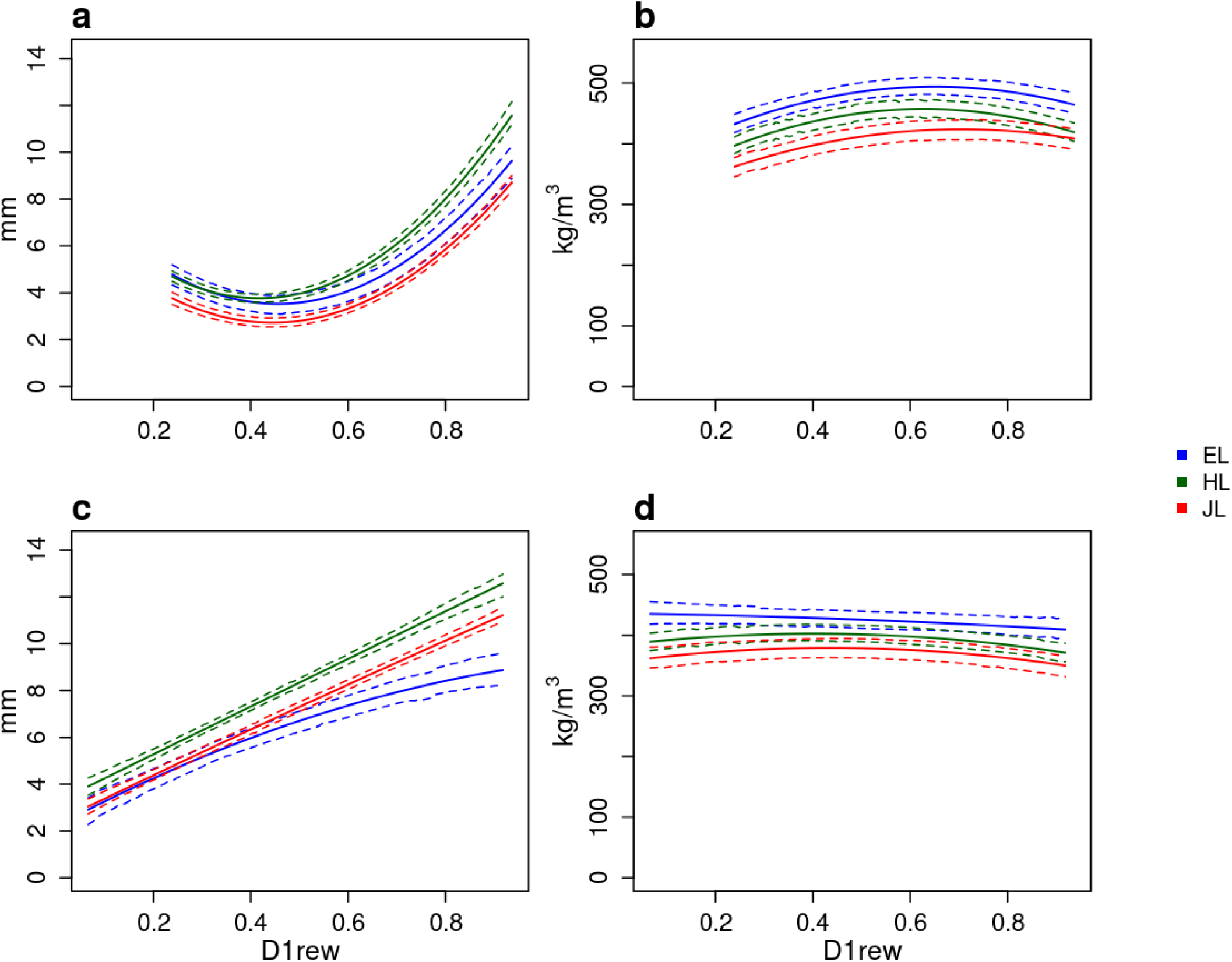
Reaction norms of ring width (a, c) and ring mean density (b, d) along the first decile of the daily relative extractable water (D1rew), in the sites SA (a-b) and SS (c-d), for each taxon: European larch (in blue, EL), Japanese larch (in red, JL) and their hybrid (in green, HL). These average reaction norms at the taxon level represent the fixed components of the order 3 random regressions. Dashed lines: 95% credible intervals

For any taxon and in any site, RW was plastic as it increased with increasing water availability (Fig. 4). The three taxa showed however different curves along the D1rew, being close to each other when water availability was minimal (D1rew close to 0), and splitting apart with increasing water availability, where the superiority of HL over its parental references was the highest. The gain in superiority for RW of HL over its parental references due to increasing D1rew ranged between +1.04 mm (HL *vs*. JL in SS) and +3.26 mm (HL *vs*. EL in SA). For high D1rew, the CIs of the HL reaction norm were not overlapping with those of the parental species.

On the opposite, the trait RMD showed neither heterosis nor plasticity. The hybrid ranged between both its parents, and all the norms of reaction were almost flat for this trait, showing no conspicuous pattern of variation along the water availability gradient D1rew.

### Heritabilities and genetic performances along the water availability gradient

All the narrow-sense heritabilities we estimated were very low (Fig. 5). Low heritabilities were overall due to high residual variances in comparison to the lower additive and permanent environment variances (Fig. 3). Despite this residual noise, we could distinguish parental performances for both traits, and some extreme performances of contrasting genotypes were different with statistical credibility (Supplementary 4, Fig. S 6-S 7).

**Figure 5.**
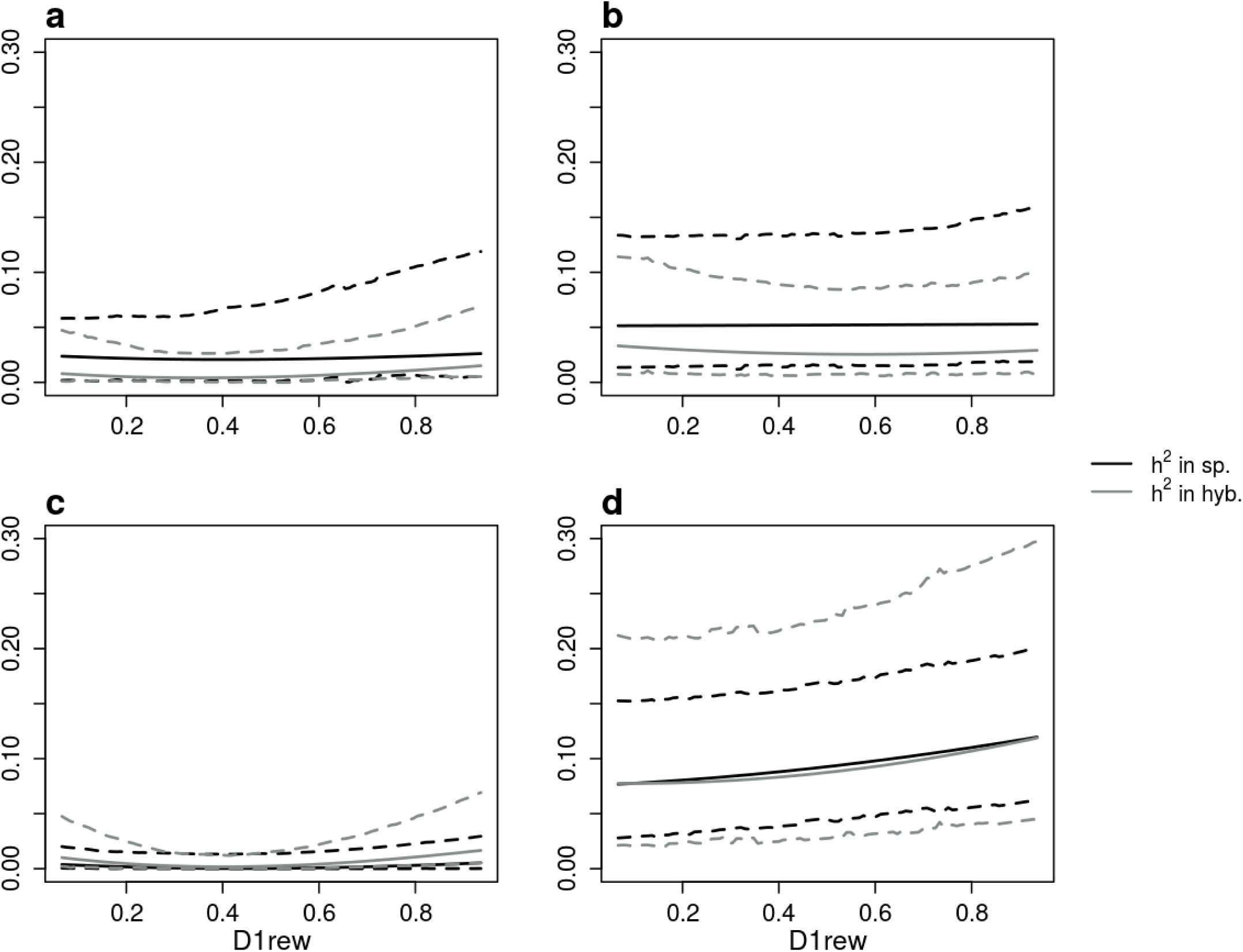
Narrow-sense heritabilities along the first decile of the daily relative extractable water (D1rew), for European larch (a-b) and Japanese larch (c-d), for the traits: ring width (a, c) and ring mean density (b, d). Black line: heritability in pure species crosses; and grey line: heritability in hybridization. Dashed lines: 95% credible intervals

Ring width heritabilities were close to 0 (Fig. 5). The signal for performance contrasts for RW in pure species was also very weak, but both species showed contrasted performances in hybridization as the water availability increased; some of these contrasts were supported by non-overlapping 95% CIs when D1rew was high (Supplementary 4, Fig. S 7).

Heritabilities for RMD were higher than those of RW, especially on the JL side for which they varied between 0.08 and 0.12 both in pure species and in hybridization (Fig. 5). Pure species heritability and heritability in hybridization were very close for JL, and they both increased with water availability. The increase in heritability for RMD on the JL side reflected an increase in the contrasts between individual performances (Supplementary 4, Fig S 6), supported by some non-overlapping 95% CIs for high D1rew (Supplementary 4, Fig S 7). Moreover, the ranking of the 9 JL parents’ performances for RMD was consistent in pure species and in hybridization (Supplementary 4, Fig S 6).

### Correlations along the water availability gradient

The additive genetic correlation between RMD and RW showed similar patterns between pure species and their respective contributions in hybridization (Fig. 6). The differences were marked when comparing EL versus JL correlation patterns across the environmental gradient. The additive correlation decreased slightly along the gradient from around 0 to −0.41 in EL pure species (−0.35 in hybridization). On the JL side, it started from a positive correlation of 0.48 in pure species (0.14 in hybridization) and it steeply switched to a negative correlation of −0.60 (−0.41 in hybridization). The change in sign occurred at D1rew between 0.4 and 0.5. Still on the JL side, 95% CI excluded 0 in pure species at high D1rew (around 0.65). Due to the higher genetic variance for RW for JL in hybridization than in pure species, the additive correlation pattern was especially visible in the JL performances in hybridization: for the highest D1rew, the ranking between RW and RMD was almost inverted (Supplementary 4, Fig S 6).

**Figure 6.**
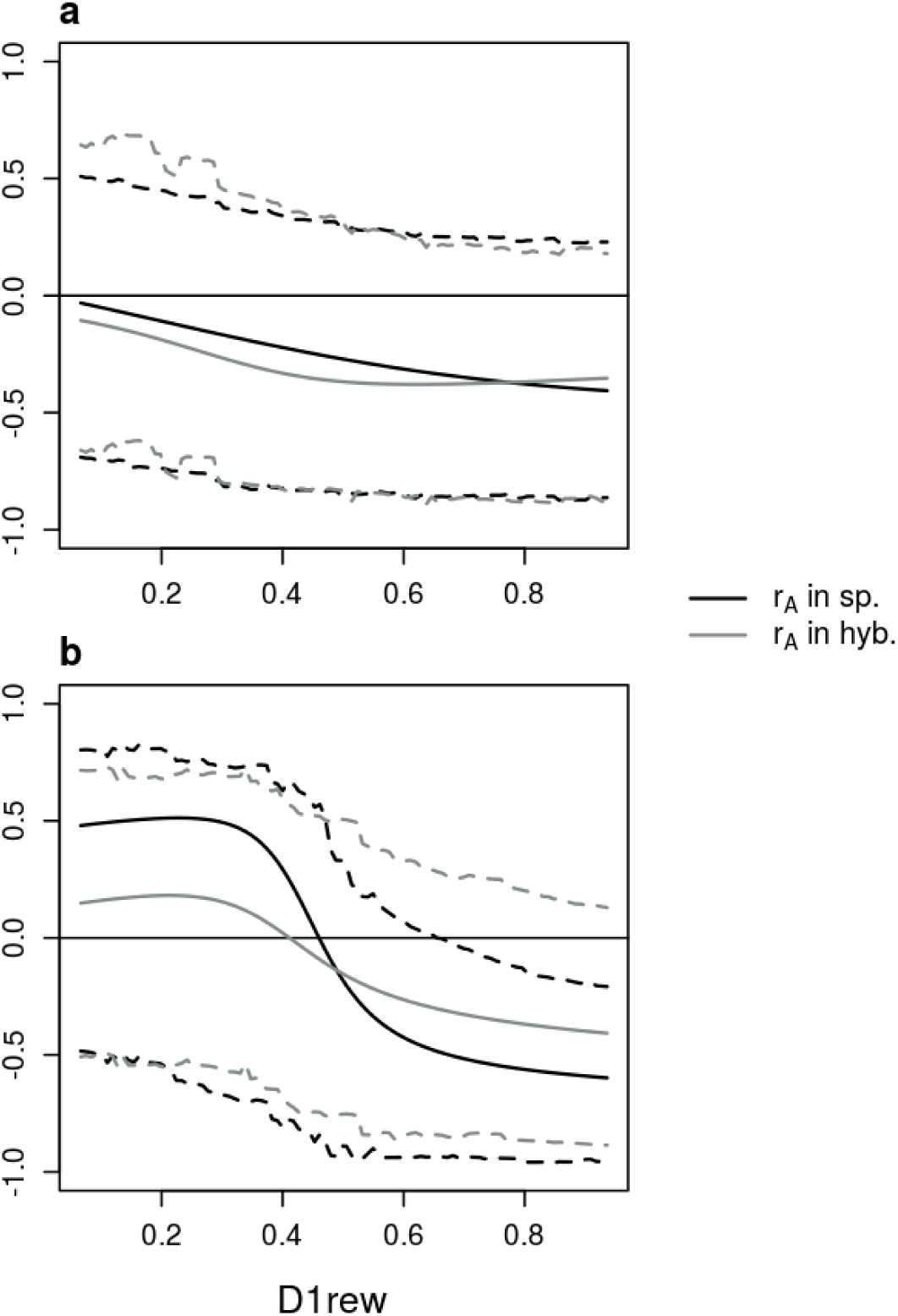
Additive correlation between ring width and ring mean density along the first decile of the daily relative extractable water (D1rew), for European larch (a) and Japanese larch (b). Black line: correlation in pure species crosses; and grey line: correlation in hybridization. Dashed lines: 95% credible intervals

The permanent environment correlation between RW and RMD was negative for HL and JL, and did not vary along the environmental gradient. It was also null to negative for EL (Supplementary 4, Fig S 8). This means that the sum of effects that the model did not explicitly account for (*i.e.* micro-environment, competition between trees, non-additive genetic effects, etc.) tended to induce a negative correlation between radial growth and wood density.

### Simulation: accuracy of the random regression model with single record per individual

Using simulated data, we evaluated the ability of the random regression model to predict the additive component of reaction norms from family series of single observations per environment. The accuracy of the model depended on the scenario and on the order of the random regression (Fig. 7). The 1^st^ scenario (*n* = 20 progenies, *h*^2^ = 0.1) showed very poor predictive abilities, independently of the order. When a larger number of progenies (*n* = 120) or a higher heritability (*h*^2^ = 0.6) was available, the accuracy was still low for order 0 (fixed and random intercept model) but it greatly increased with order 1 (addition of fixed and random slopes). However, in each case (*n* = 120 or *h*^2^ = 0.6) no further accuracy was gained from order 1 to order 2. Only the accuracy for the last scenario (both *n* = 120 and *h*^2^ = 0.6) increased with order 2 (addition of fixed and random parabolas). The accuracy for the last scenario analyzed with order 2 model ranged between 0.905 and 1 depending on the simulation and on the position along the environmental gradient.

**Figure 7.**
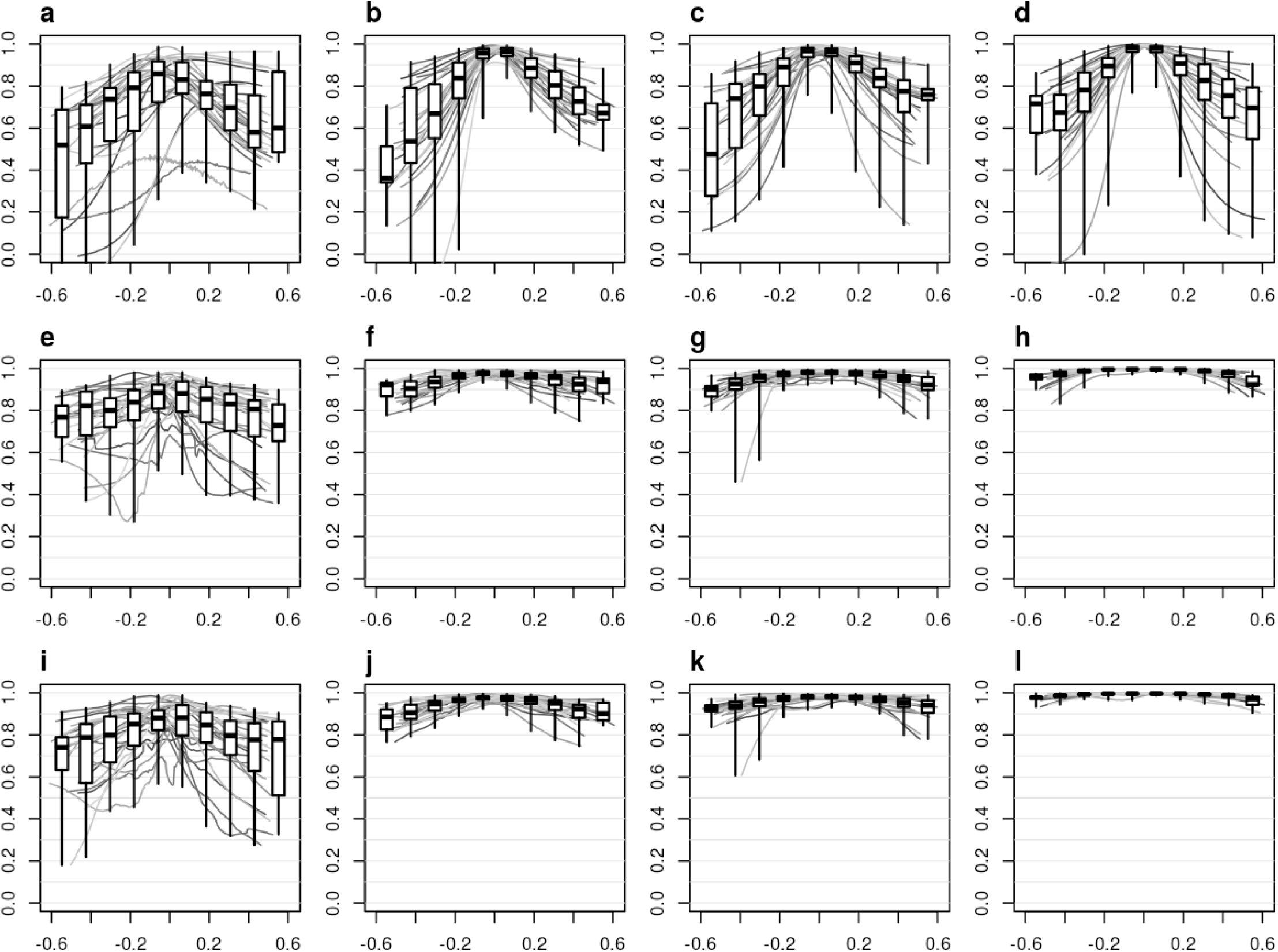
Accuracy of the predictions of parents’ additive reaction norms in each scenario: (1) *h*^2^ = 0.1 and *n* = 20 (a, e, i), (2) *h*^2^ = 0.1 and *n* = 120 (b, f, j), (3) *h*^2^ = 0.6 and *n* = 20 (c, g, k), and (4) *h*^2^ = 0.6 and *n* = 120 (d, h, l); for 100 simulations in each scenario, and for each order of random regression: order 0 (a-d), order 1 (e-h) and order 2 (i-l). Each grey curve is the accuracy of 1 simulation; boxplots summarize all the point accuracies in segments of a 10^th^ of the simulated environmental gradient

The order 0 models were not able to estimate the additive variance properly (Supplementary 4, Fig. S 9). From order 1 and over, the ability of the model to estimate the additive variances appeared overall dependent on the heritability. Indeed, the estimated additive variance were close to the true ones in scenarios 3 and 4 (both *h*^2^ = 0.6) with order 1 or 2. With lower heritability (*h*^2^ = 0.1) and order 1 or 2, the additive variance were overestimated.

## DISCUSSION

In this study, we constructed the reaction norms of annual wood-formation traits along a water availability gradient in a larch multi-site diallel mating experiment. The reaction norms were fitted using random regression modeling, allowing to assess the changes in heritability and genetic correlations along the gradient. Our study was complemented by using simulations involving the same analytical approach, where we evaluated the ability of the random regression model to estimate the additive component in a frequent phenotypic plasticity experimental setting: that of parental reaction norms from non-repeated observations of their progenies as data.

The annual ring width was plastic and as expected increasing water availability allowed a higher radial growth (Fig. 4). At the taxon level, the hybrids performed better than their parents in each of the two sites of the study. This alternative modeling confirmed our previous findings (Marchal *et al.* 2017), in that hybrid larch had a stable superiority across sites. However, within each site, HL demonstrated more plasticity than its parental references: indeed, under water stress all taxa produced a similarly narrow ring, whereas in favorable conditions of higher water availability the HL expressed superiority over its parental references. In other words, the HL reaction norms were steeper than those of the parental counterparts over the same gradient.

On the contrary, the second trait under study, RMD, was not plastic for this water availability gradient and expressed no heterosis in any site. Although RMD displayed higher narrow-sense heritabilities (*h*^2^) than RW, it did not reach high values, with a global maximum of only 0.12. Despite the weak genetic signal, the performance functions of the 18 parents of the diallel could be represented distinctively across the water availability gradient, as well as the functions corresponding to the genetic parameters (*h*^2^ and *r_A_*, the additive correlation between RW and RMD). Interestingly, on the Japanese larch side, *r_A_* switched from positive to negative values as water availability increased, with the highest negative values being different from zero with statistical credibility (Fig. 6). The emergence of this negative additive correlation might be explained by an increase in the early wood / late wood ratio with increasing water availability, with early (spring) wood being generally less dense than that of late (summer) wood (Fig. 1). However, we need to look more carefully to other ring traits (as did, *e.g.*, Bryukhanova and Fonti 2013) and their respective correlations before proposing any causal explanation, with the goal to better understand the structure of the genetic variability of larch wood plasticity.

We obtained fairly high *R*^2^ for the reaction norms models, suggesting that water plays an important role in the tree ring phenotypic plasticity. We evaluated a simpler environmental factor (sum of daily rain minus potential evapotranspiration from May to August) but it showed a lower *R*^2^, highlighting the relevance of our water availability index ‘D1rew’. It should be noted, however, that our water balance model has not been field-calibrated, and it should then be considered with care if generalizations are to be made. Moreover, the index D1rew gives no indication on the distribution of the driest days along the year. The timing of a water deficit, in spring or in summer, could have more or less effect on different ring traits; for instance, those relative to early or late wood, or to the transition between the two. It also has to be said that the relation between water balance and radial growth that we showed in the present study does not necessarily imply a direct causality. Indeed, other factors may play a role in the observed plastic response. For instance, heat affects directly the photosynthetic efficiency and the resources that may be allocated to growth (Rennenberg *et al.* 2006). Heat and drought being highly correlated, their effects could well be confounded to some extent. More broadly, the environment has a multivariate nature. Soil, climate, but also competition with neighboring trees are known to affect the tree’s growth. We isolated what we expected to be one of the most important environmental factor for radial growth, yet the existence of an important site effect pinpoints the fact that some other environmental factors might be involved in the tree ring phenotypic plasticity. Identifying relevant environmental factors of plant plastic reactions is an open area of research, notably in the context of global warming. The present study did not aim explicitly at the identification of relevant environmental triggers, rather it presented an approach that could help in such identification.

We studied phenotypic plasticity at two levels. The first level was spatial, at the across-site scale, and the second level was longitudinal, at the individual scale. At the across-site scale, heterosis was shown to be stable, supporting the common statement that hybrids are more stable across macro-environmental sites than their parental counterparts (Gallais 2009). Specifically, the ranking of the parents species varied across sites whereas hybrid was invariably the highest performing taxon, in what could be qualified as hybrid homeostasis according to the theory developed by Knight (1973). The spatial plasticity is generally, and historically, the one that interests breeders the most because of the operational implications for the deployment of varieties. Using non-linear random regressions, Marcatti et al. (2017) fitted eucalyptus growth reaction norms along a gradient of spatial environments. The spatially distributed climatic environment was described with principal component analysis to account for its multivariate nature. This method is very appealing in order to deal with spatial phenotypic plasticity in tree breeding. However, Marcatti *et al.* (2017)’s approach could be further improved by accounting for pedigree information such as a mating design, from which quantitative genetic parameters could be estimated (Hinkelmann 1974; Falconer and Mackay 1996). In this context, the use of random regressions as defined by Kirkpatrick and Heckman (1989) and presented in this paper, or other covariance functions, would be relevant.

The second level of phenotypic plasticity that we addressed was longitudinal, along the year-to-year water availability gradient. The resulting reaction norms were strongly influenced by the sites. Indeed, our study steps in the direction that tree ring longitudinal phenotypic plasticity should be seen as a plastic trait in itself, varying spatially (De Luis *et al.* 2013; Natalini *et al.* 2016), varying with long-term trends such as global warming (Natalini *et al.* 2016), and varying with the level of competition between neighboring trees in wet years (Carnwath and Nelson 2016). Growth recovery, the ability for trees to produce large rings the years following a drought event, is also site-dependent (Gazol *et al.* 2017). Besides the site effect, a substantial part of the individual variation was shown to occur between taxa, with hybrid larch being more plastic and at a higher mean level than its parental counterparts. Therefore, this increase in longitudinal plasticity appears as a broader picture of the heterosis that was observed on an integrative scale across years, namely in the total circumference (Marchal *et al.* 2017).

The molecular and physiological mechanisms behind the increased plasticity of hybrid larch remain open questions. Longitudinal plasticity as shown here is still a novel approach, with less immediate application to current plant breeding, unlike spatial plasticity. It certainly opens up new possibilities whenever long-time series are available, where extreme events, such as the 2003 drought in France (Bréda *et al.* 2006; Rennenberg *et al.* 2006), are recorded. Some initiatives, for instance, compared dead trees *vs.* alive neighbors immediately after extreme climatic events for their past wood records (Martinez-Meier *et al.* 2008), finding that both classes had long-term distinctive patterns of reaction. This kind of study could well be undertaken with a random regression approach to gain insight in the quantitative genetics of such longitudinal patterns and their environmental drivers, and be of potential use ultimately for breeders.

One eventual problem with longitudinal data is autocorrelation. In order to minimize its effects, several authors propose an extra step that consists in fitting an autocorrelation model on the tree ring series. The resulting residuals, that are more independent than the raw data, are then used as the response variable in the subsequent phenotypic plasticity models (Bryukhanova and Fonti 2013; De Luis *et al.* 2013). In this study, we did not do so because the chronology was somehow already involved in the environmental variable. Indeed, as the trees grew older, the stand LAI increased, and so did the transpiration, making water generally less available. Though this trend was not so strong (rain and potential evapo-transpiration during the growing season were the main drivers of D1rew, as seen on Supplementary 1, Fig. S 2), we did not want to account twice for the same chronological effect, and therefore we chose to work with raw data instead. The much weaker explicative power of D1rew of the previous year compared to that of D1rew of the current year supports our decision. We acknowledge though that there remains a risk of non-controlled autocorrelation in the data, in particular due to the trees’ ontogeny and to the onset of competition between trees (Sánchez *et al.* 2013).

The simulations showed that it was possible to estimate the additive component of reaction norms using only singular observations of related individuals. However, the quantity and the quality of information (*i.e.* respectively, the number of related individuals and the heritability) were key factors to estimate properly the additive components of reaction norms. Although the simulation was not meant to mimic the real case in the genomic layout of effects, it pinpointed the eventuality of potential biases in the estimation of the genetic variances (Supplementary 4, Fig. S 9). As emphasized by Misztal *et al.* (2000), a limitation of the model used in the present study is the lack of covariance function for the residuals. This limitation could be a source of bias in the estimation of variance components and of heritabilities. Unfortunately, this feature was not available yet with the software we used. Indeed, we fitted linear mixed models in which the covariance functions were implicit and computed from the covariance between the regression coefficients. On the other hand, in the simulation, the additive reaction norms were properly estimated.

Random regression is already used for the modeling of reaction norms, especially in dairy cattle for which industry produces a large flow of longitudinal data (*e.g.* Kolmodin *et al.* 2002; Windig *et al.* 2006; Santana *et al.* 2017). However, plasticity has been suggested (Bradshaw 1965) and demonstrated (Murren *et al.* 2014) to be of special importance in plants. Indeed, because they are sessile, plants have to face their environment in a different way than animals that are capable of behavioral responses and locomotion. Manifestations of phenotypic plasticity have been reported in several perennial crops. For instance, grape vine manifests phenotypic plasticity in terms of fruit weight and chemical composition (Dai *et al.* 2011). Even in equatorial regions, oil palm is able to react to subtle variations in photoperiod and drought events by changing its bunch productivity (Legros *et al.* 2009). Cherry tree phenology reacts promptly to climate, notably heat, with global warming expected to bring flowering a month forward (Allen *et al.* 2014). Ismaili *et al.* (2016) showed a significant genotype-by-year interaction in apricot tree, and recommend the use of mixed models for the analysis of perennial plants’ longitudinal data. All these examples could be good candidates for analysis based on covariance functions. Indeed, the possibility to define quantitative genetic parameters as functions of the environment and to model the additive contributions to phenotypic plasticity opens wide perspectives in terms of selection (De Jong and Bijma 2002), especially in a global warming context (Koski 1996).

Like trees, other organisms naturally cumulate growth records that reflect their reaction to past environmental conditions, etched in hard organs that grow incrementally: for instance, fish otoliths, mollusc shells, corals, whale ear plugs, ibex horns, *etc.* (reviewed by Morrongiello and Thresher 2015). Morrongiello and Thresher (2015) advocate for the use of random regression for the analysis of such natural records of longitudinal data, especially with regards to these species’ phenotypic plasticity. The random regression framework as developed by Kirkpatrick and Heckman (1989) exhibits two particular strengths that deserve to be emphasized once more. First, the use of orthogonal base functions, such as Legendre polynomials, allows the fit of virtually any shape of growth curves or reaction norms. Although the example we presented here did not illustrate this need, neglecting curvature when studying evolutionary divergence in reaction norms leads to a risk of missing critically important information (Murren *et al.* 2014). The traditional use of *x* and *x*^2^ as covariates should be avoided in any case, given the high correlation that binds the identity and the square (and any power) functions.

A second point of interest is the fact that genetic information can be taken into consideration in the model, which can be of relevance not only for breeding but also in ecology studies looking for drivers and patterns of natural selection (Brommer *et al.* 2005). As we illustrated with the results of our simulations, this possibility opens up the use of a random regression framework to species whose individuals do not cumulate in any known form longitudinal records of their plastic responses. In this sense, our simulation provided an example of random regression being an alternative to traditional methods: related individuals can indeed give access to the additive component of the reaction norms, and this can likely be extrapolated to isofemale lines (Gibert *et al.* 2004) or half-sib families (Valladares *et al.* 2006). Finally, the pedigree information can be conveniently replaced by molecular information (*e.g.* Ly *et al.* 2018), extending the potential of the random regression framework beyond the limitation of our capability to realize time-consuming, sometimes impossible, artificial mating.

## Acknowledgements

The authors sincerely acknowledge the technical staff of INRA experimental units (UE GBFOR and UE BIOGECO) who have established, maintained and assessed the field trials as well as collected the increments cores. The authors thank Genobois platform technical staff who have prepared and X-rayed wood samples and provided microdensitometry profiles. We thank Philippe Rozenberg for his implication all along the paper. We thank the think-tank ‘PlasPhen’ for providing opportunities to move forward in our thinking on this paper, and in particular Patricia Gibert and Vincent Debat who organized it. We thank Isabelle Cousin and Ghislain Girot for their help with the interpretation of the soil data. We thank Vincent Ducrocq for his advices with the random regression model. We thank Météo-France and the platform INRA CLIMATIK for the availability of the climatic data. F. Muñoz was partially funded by research grant MTM2016-77501-P from the Spanish Ministry of Economy and Competitiveness. F. Muñoz and L. Sánchez received funding from the European Union’s Seventh Framework Program for research, technological development, and demonstration under grant agreement no. 284181 (‘Trees4Future’, coord. L.E.Pâques).

---

**Figure S 1.**
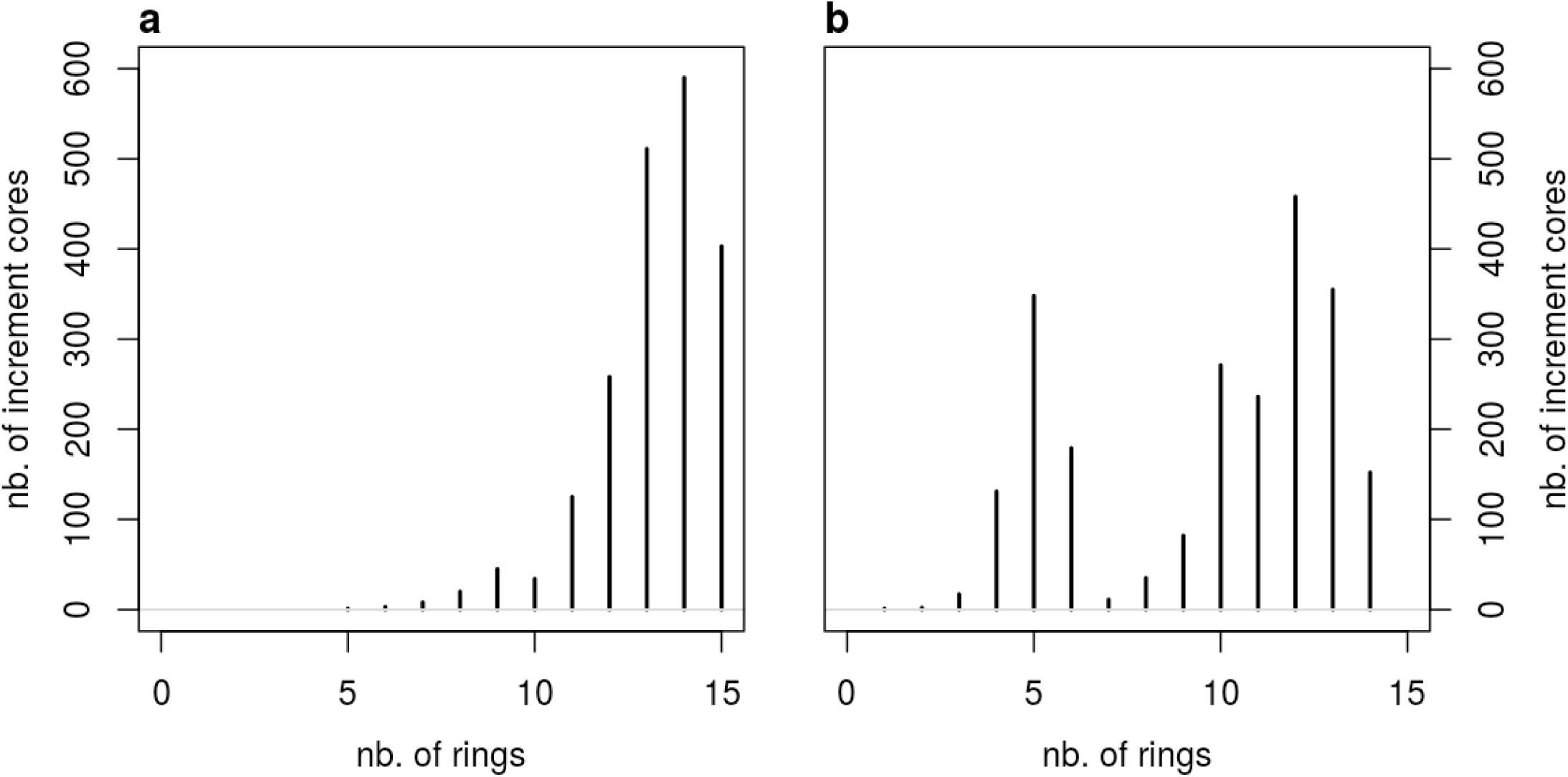
Number of rings per increment cores from Saint-Appolinaire (a) and from Saint-Saud (b)

**Figure S 2.**
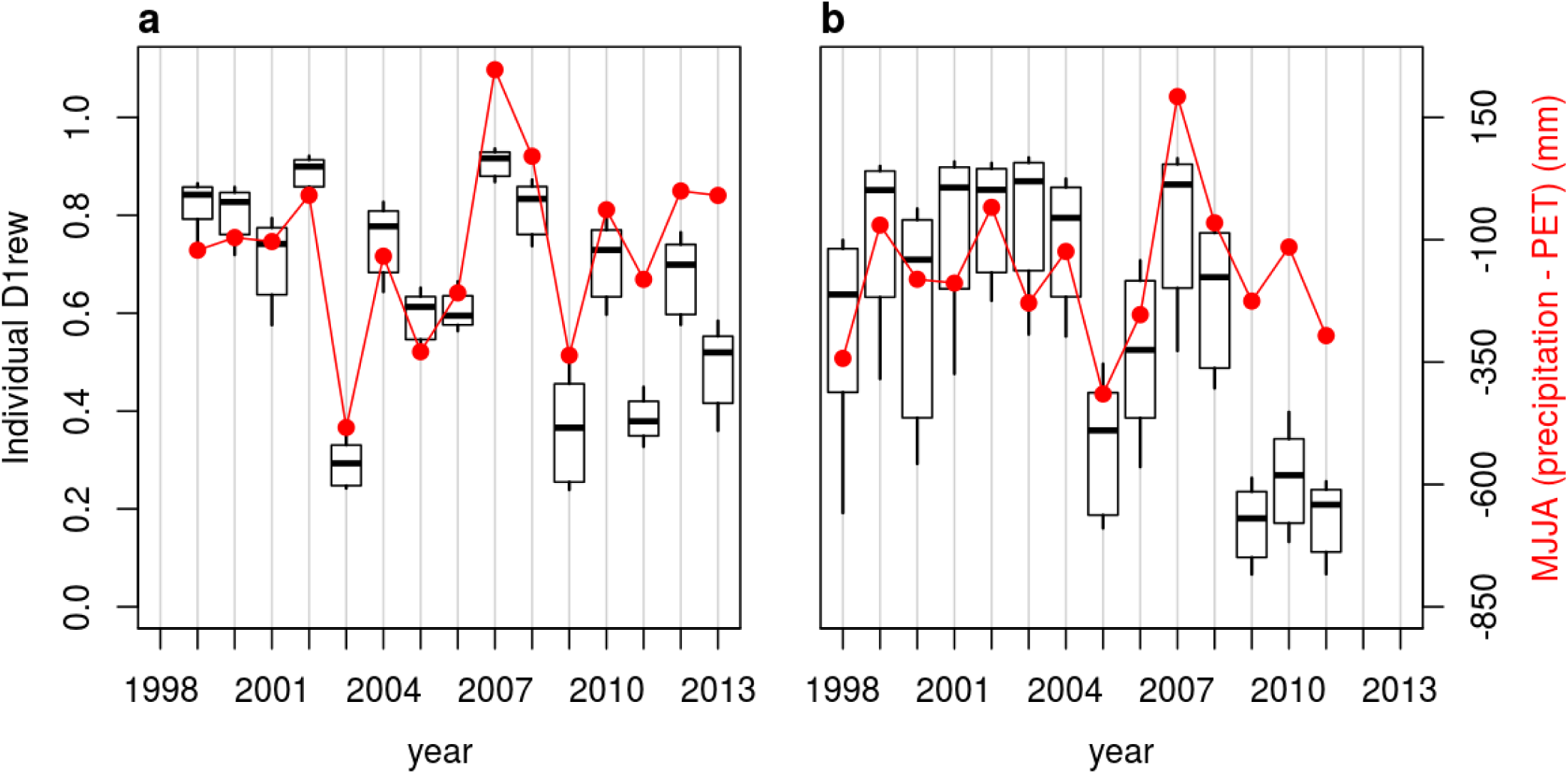
In black: boxplots of the distribution of individual D1rew (first decile of the daily relative extractable water) for each year in each site SA (a) and SS (b). In red: the simpler index MJJA, defined as the sum of the daily differences between precipitation and potential evapotranspiration from May to July

## SUPPLEMENTARY MATERIAL 2 WATER BALANCE MODEL

### Leaf area index

The leaf area index (LAI) is the ratio of leave surface per ground surface. Thus in a growing stand the LAI is expected to increase, and this plays an important role in the water balance model, as detailed further. The LAI can be calculated from the basal area, that is, the surface of cross-section of tree stems per ground surface. Basal area was calculated in each site at each age for which we had a breast-height circumference (BHC) measurement. We estimated transmittance from basal area and age using Sonohat *et al.* (2004) (‘Model 2’, *R*^2^ = 0.867):

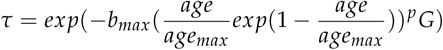

where *G* was the basal area, and *age_max_*, *p*, and *b_max_* the model parameters available in Sonohat *et al.* (2004). From transmittance we could estimate the LAI as LAI_*max*_ = −*ln*(*τ*)/*k* with the value *k* = 0.6 for larch (Takeda *et al.* 2008). The index *’max’* in LAI_*max*_ means: the LAI when the vegetation is maximal in the season. Then, we linearly inferred the LAI_*max*_ at any age for which we had ring observations. The LAI_*max*_ was inferred at the site scale. We present the calculated and the inferred LAI_*max*_ in Fig. S 3.

**Figure S 3.**
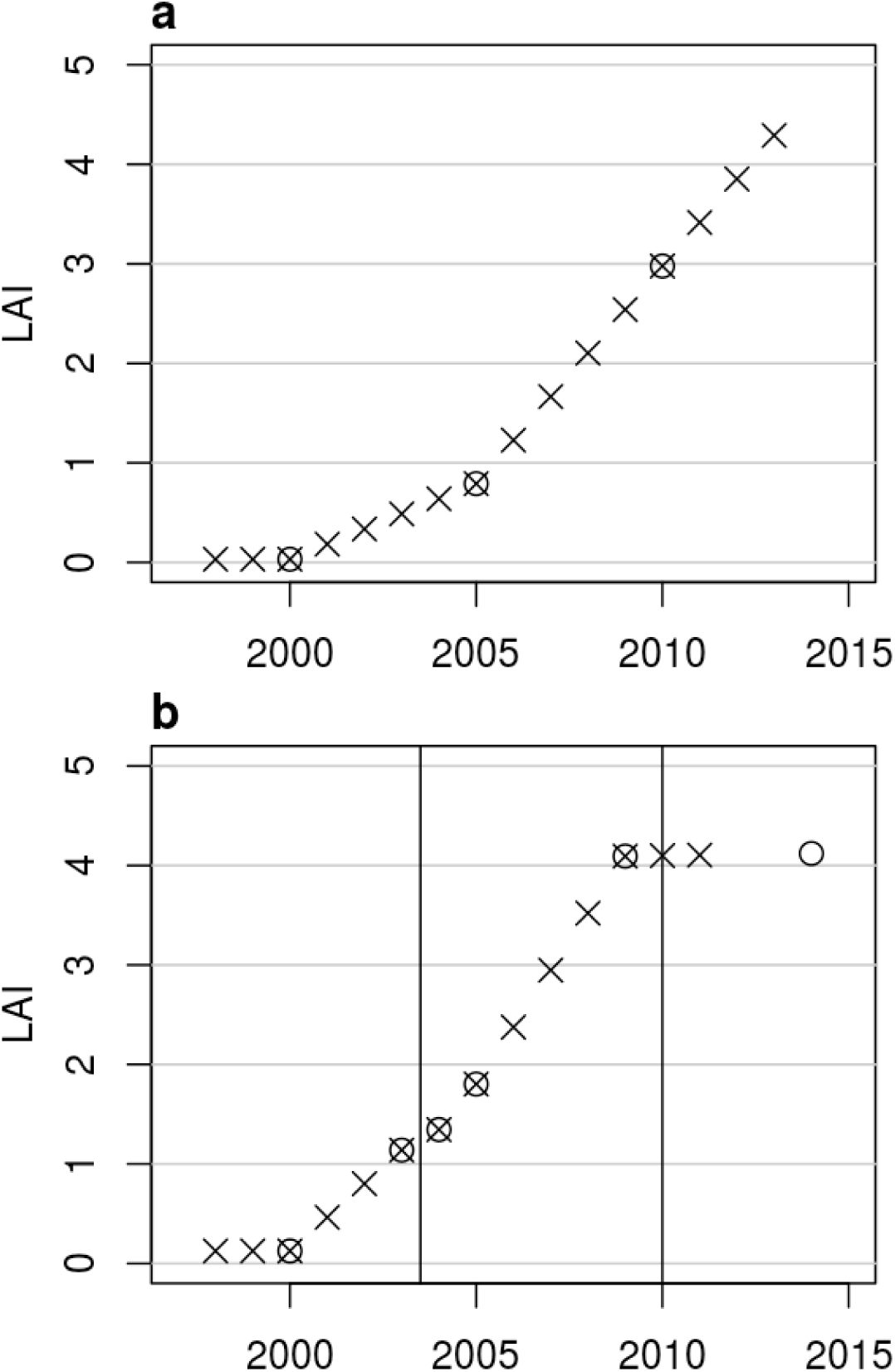
LAI_*max*_ evolution in each site: SA (a) and SS (b). Circles: LAI_*max*_ estimated from basal area. Non-circled crosses: LAI_*max*_ linearly inferred. All crosses: LAI_*max*_ used in the water balance model

### Soil available water capacity

We excavated 2 pits in SA and 3 pits in SS, in the most contrasted areas. The contrasted areas were assessed using tree height (at the last age available) spatial effect maps. The spatial effects were predicted as best linear unbiased predictions (BLUPs) from model ‘M1’ in Marchal *et al.* (2017). In each pit, we measured soil horizon thickness, stone content, and we collected samples to assess the soil texture. We estimated the available water capacity (AWC) for each pit (Bruand *et al.* 2004), that is, the maximal amount of water available for plants that the soil can store. We assumed that height spatial effect informed on AWC, as empirically supported in Fig. S 4. Within each site, we considered a linear relation between tree height spatial effects (BLUPs) and AWC in order to infer AWC at the tree level. Nevertheless, for each site, we prevented the individual trees AWC from being lower (or higher) than the lowest (or highest) pits’ AWCs, resulting in an inverse-M shaped distribution.

**Figure S 4.**
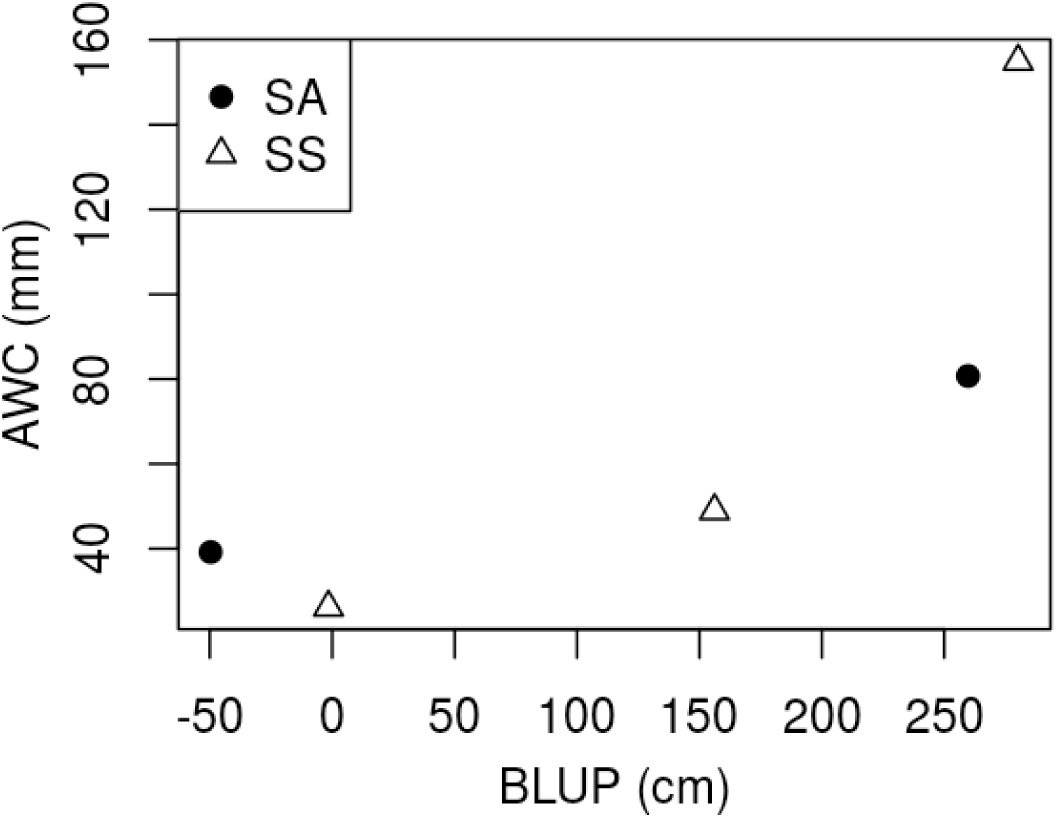
Link between height spatial effect (estimated as a best linear unbiased predictor, BLUP) and the available water capacity (AWC) in each site: Saint-Appolinaire (SA, black dots) and Saint-Saud (SS, white triangles). Each dot represents a pit

### Climatic data

The daily climate information we used were the precipitations (*P*) and the potential evapotranspiration (*PET*). Raw climatic data were from Météo-France, via the platform INRA CLIMATIK which computed the PET with Penman-Montheith method. Ten PET data points were missing in SS, we imputed them using an autoregressive model (?) with an autocorrelation parameter *ρ* = 0.95.

### Water balance model and water availability indexes

We used an adapted, simplified implementation of Granier *et al.* (1999)’s daily water balance model. We simplified the model as follows: (i) we proposed a simplified formulation for the understorey evapotranspiration; (ii) we proposed a simplified formulation for the rainfall interception; (iii) we ignored soil stratification and distribution of the roots, so only the overall AWC described the subsoil. The daily water balance was:

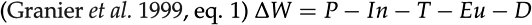

with *W*: the soil water content and Δ*W* its daily variation; *P*: the precipitation; *In*: the rainfall interception; *T*: the overstorey transpiration; *Eu*: the understorey evapotranspiration; and *D*: the drainage; all expressed in mm. The relative extractable water (REW) content in soil was calculated as REW = *W*/AWC. The LAI was 0 until day 105 (15^th^ of April for a non-leap year), then increased linearly in 30 days, stayed at LAI_*max*_ a while and finally decreased linearly to 0 in 30 days, finishing the decrease at day 288 (15^th^ of October). The canopy transpiration *T* was calculated as following:

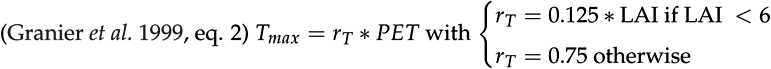

If REW was above 0.4 (considered as a drought threshold), *T* was *T_max_.* Otherwise, *T* decreased linearly with REW. We used no intercept to the linear relation between *T* and REW, so that the transpiration was null if no water was available. The drainage *D* was such as REW was never above 1. We proposed the following method for Eu:

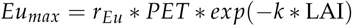

with *k* = 0.6 for larch (Takeda *et al.* 2008); then *Eu* was calculated from *Eu_max_* the same way *T* was calculated from *T_max_*. Using a model derived from Penman-Montheith equations, Kelliher *et al.* (1995) represented the relation between *Gs/Gc* and the LAI; with *Gs* the understorey surface conductance and *Gc* the tree canopy conductance. This relation is plotted in Fig. S 5. We also present in Fig. S 5 the ratio (*Eu* + *T*)/*T* depending on *r_Eu_*. It arises from this comparison that *r_Eu_* = 0.375 provides water flow ratios that are consistent with the conductance ratios from Kelliher *et al.* (1995), especially in the condition of low LAI. Therefore, we used *r_Eu_* = 0.375 to parameterize the model during the growth season. In order to account for the decrease in the overstorey biological activity in winter, *r_Eu_* was set to 0.125 between the days 288 and 105 with 30 days of linear transition (just as the trees LAI varying from LAI_*max*_ to 0).

**Figure S 5.**
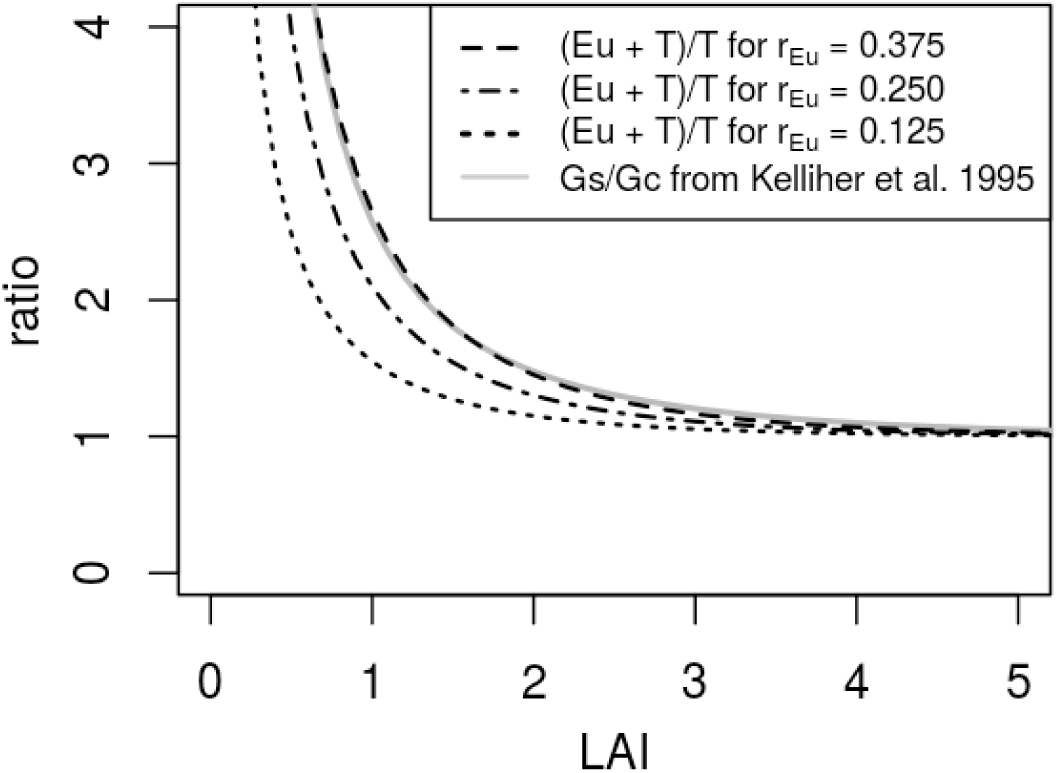
Ratio between the water transpired by tree canopy (*T*) and evapotranspired by the understorey vegetation (*Eu*), and ratio between understorey vegetation conductance (*Gs*) and tree canopy conductance (*Gc*) as modeled by Kelliher *et al.* (1995), depending on the leaf area index (LAI)

In order to model the water losses by rainfall interception we introduced a new compartment, the tree canopy water storage *S.* This compartment was limited by *S_max_*, the total amount of water that the canopy could store, which was set as a function of the LAI: *S_max_* = 3 mm * (1 − *exp*(−*k* * LAI_*max*_)). The value 3 mm and the absence of seasonal variation for the canopy water storage capacity were specific to larch (Reynolds and Henderson 1967). The interception algorithm was unsophisticated: S was filled first, then when it was full the extra water dropped to the ground. We applied the following rules adapted from Granier *et al.* (1999): (i) The canopy transpiration *T* was reduced by 20% of *S* and (ii) the sum of *T*, *Eu* and the evaporation from *S* was limited to 1.2 times the *PET.* Finally, the remaining water on the leaves was transferred to the next day.

### SUPPLEMENTARY MATERIAL 3 SIMULATOR

A locus-based simulation software (Metagene) using a finite loci approach was previously developed at INRA (Sánchez *et al.* 2008, http://www.igv.fi.cnr.it/noveltree). The initial software was adapted to the study of forest tree breeding strategies dealing with adverse genetic correlations (Hallingbäck *et al.* 2014), or long-term genetic diversity issues (Wu *et al.* 2016). It was further expanded for the present study to simulate heritable traits responding longitudinally to environmental variations. According to the phenotypic plasticity literature (Windig *et al.* 2004; Pigliucci 2005), two nonexclusive genetic mechanisms can be assumed for modeling a plastic response: ‘epistatic’ plasticity and ‘pleiotropic’ plasticity. Briefly, while epistatic plasticity denotes mainly the causal mechanism by which regulatory genes serve as environmentally operated signal boxes for switching between alternative genetic pathways, the pleiotropic plasticity concept refers to genes that have pleiotropic effects on a given character expressed in different environments. Both mechanisms were coded as available gene actions in the simulator, but only the latter was used in this study for simplicity. Thus, the effect of a locus was set as a function of a given environmental gradient x. Rather than prospecting the effects of underlying genetic architectures and mechanisms on the plastic response, our main objective here was to produce with a reasonably simple setup heritable reaction norms.

Therefore, our modeling of plastic responses relied on loci with alleles showing environmental sensibility in their genic effects. For this, genotypic values were modeled by a quadratic function such as *α*_(_*x*) = *α*_0_ + *α*_1_ (*x* + *δ*) + *α*_2_(*x* + *δ*)^2^, where parameters *α*_0_, *α*_1_, *α*_2_ and *δ* defined a genotypic reaction norm with a certain parabolic shape over the range of the environment x. Each locus required two of these functions, one for the favorable homozygote (AA) and another for the unfavorable homozygote (aa), with heterozygote (Aa) being always intermediate (*i.e.* no dominance). Averaged allele effects were further computed following Falconer and Mackay (1996). Underlying the plastic trait, we considered 30 diallelic loci with alternating quadratic functions across the genome between two possible parametric sets (first set, AA: *α*_0_ = 0, *α*_1_ = 2, *α*_2_ = 0; aa: *α*_0_ = 0, *α*_1_ = −1, *α*_2_ = 0; and second set, AA: *α*_0_ = 0, *α*_1_ = 0, *α*_2_ = 2; aa: *α*_0_ = 0, *α*_1_ = 0, *α*_2_ = −3). The *δ* parameter was locus specific and varied between −0.7 and 0.75 across the genome, in such a way that a certain level of heterogeneity in *α*_(_*x*) when *x* → 0 was produced. The environmental deviate, *x*, was randomly sampled from a normal distribution (mean = 0 and standard deviation = 0.2) to obtain a set of *n* environments, equal to the number of sibs per mating.

All loci were considered to be evenly spaced across the genome, and recombination occurred without interference considering an arbitrary genome size of 600 cM. The initial sample of alleles that made up founder genotypes was randomly drawn from a distribution where allelic frequencies could be set randomly across loci within a range between 0.2 and 0.8. Individual genotypic values per environment were the result of the sum of all loci’s genotypic contributions for the corresponding environment. The corresponding phenotypic value, expressed in a given environment x, was the sum of the genotypic value and a residual deviation which was sampled from N(0, 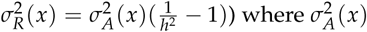 was the additive variance at the *x* environment, and *h*^2^ the initial narrow-sense heritability being constant along the environmental gradient. Note that the residual deviation was not correlated to the environmental cue *x*.

**Figure S 6.**
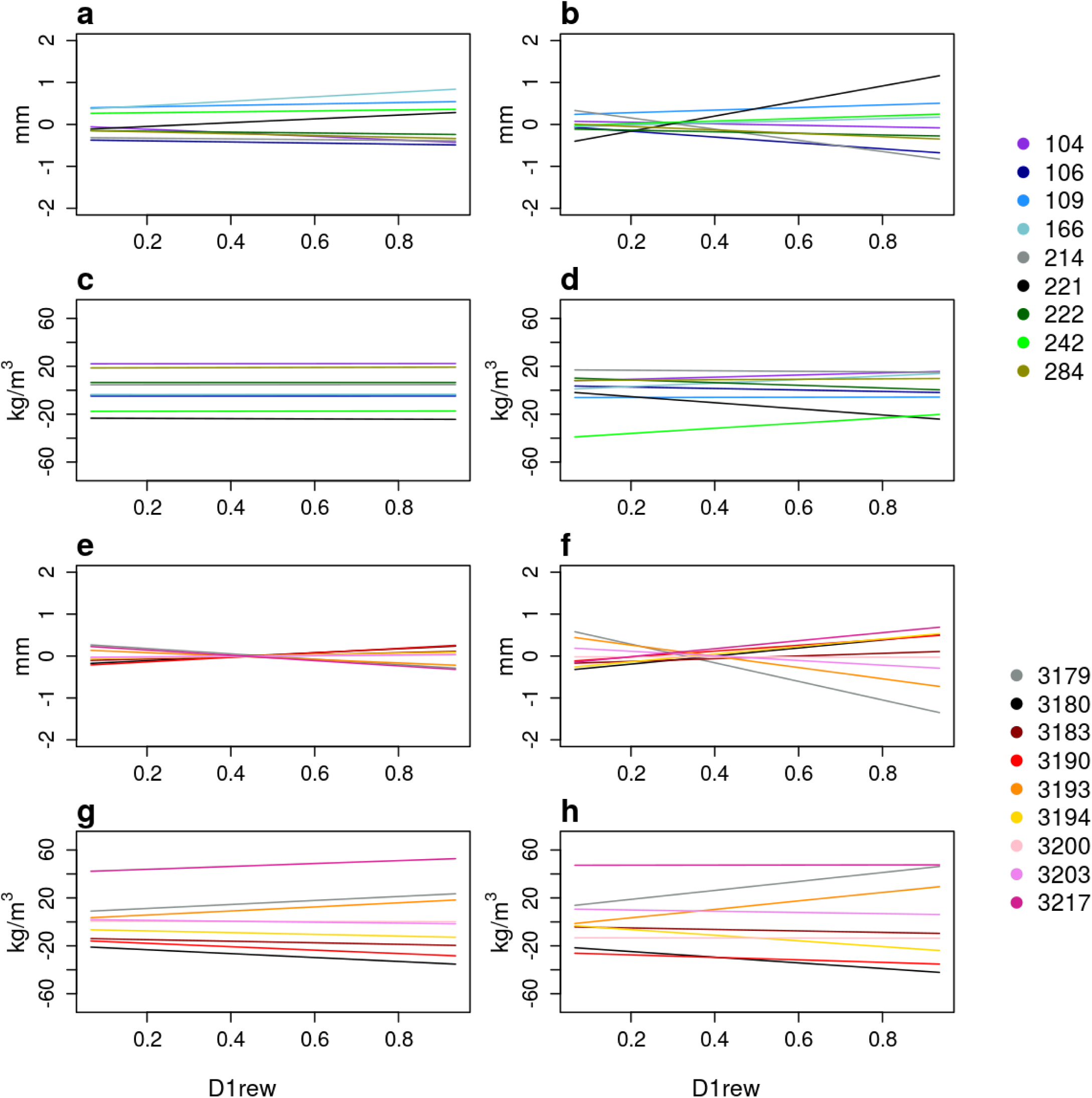
Genetic performances for the 9 European larch parents (a-d) and the 9 Japanese larch parents (e-h) for ring width (a-b, e-f) and ring mean density (c-d, g-h), in pure species (breeding value) (a, c, e, g) and in hybridization (twice the general hybridization ability) (b, d, f, h), along the first decile of the daily relative extractable water (D1rew). Each color represents one genotype, labeled in the legend on the right

**Figure S 7.**
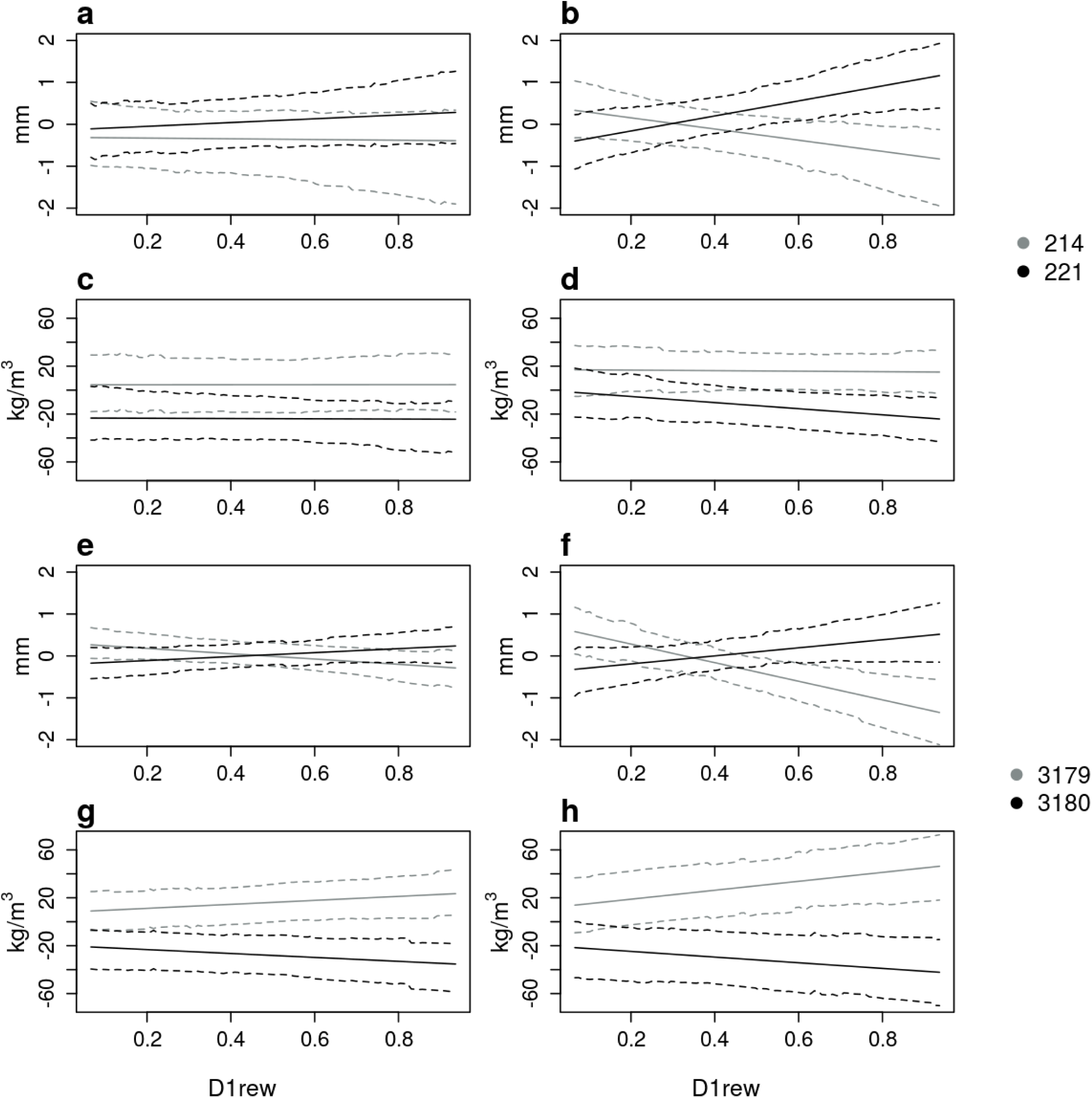
Genetic performances with 95% CIs (dashed lines) for some contrasted European larch parents (a-d) and Japanese larch parents (e-h) for ring width (a-b, e-f) and ring mean density (c-d, g-h), in pure species (breeding value) (a, c, e, g) and in hybridization (twice the general hybridization ability) (b, d, f, h), along the first decile of the daily relative extractable water (D1rew). Each color represents one genotype, labeled in the legend on the right

**Figure S 8.**
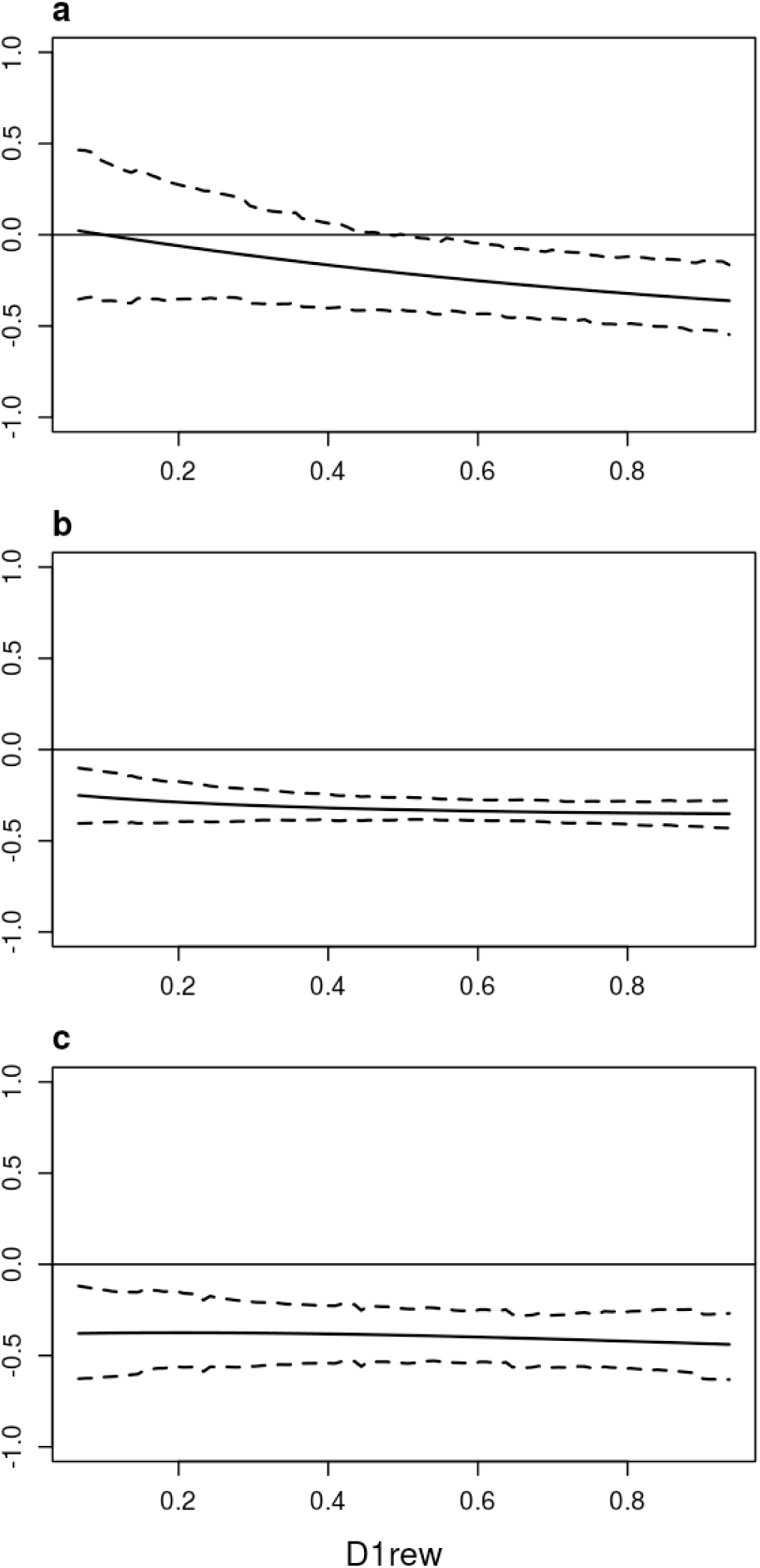
Permanent environment correlations between ring width and ring mean density along the first decile of the daily relative extractable water (D1rew), for European larch (a), hybrid larch (b) and Japanese larch (c). Dashed lines: 95% CIs

**Figure S 9.**
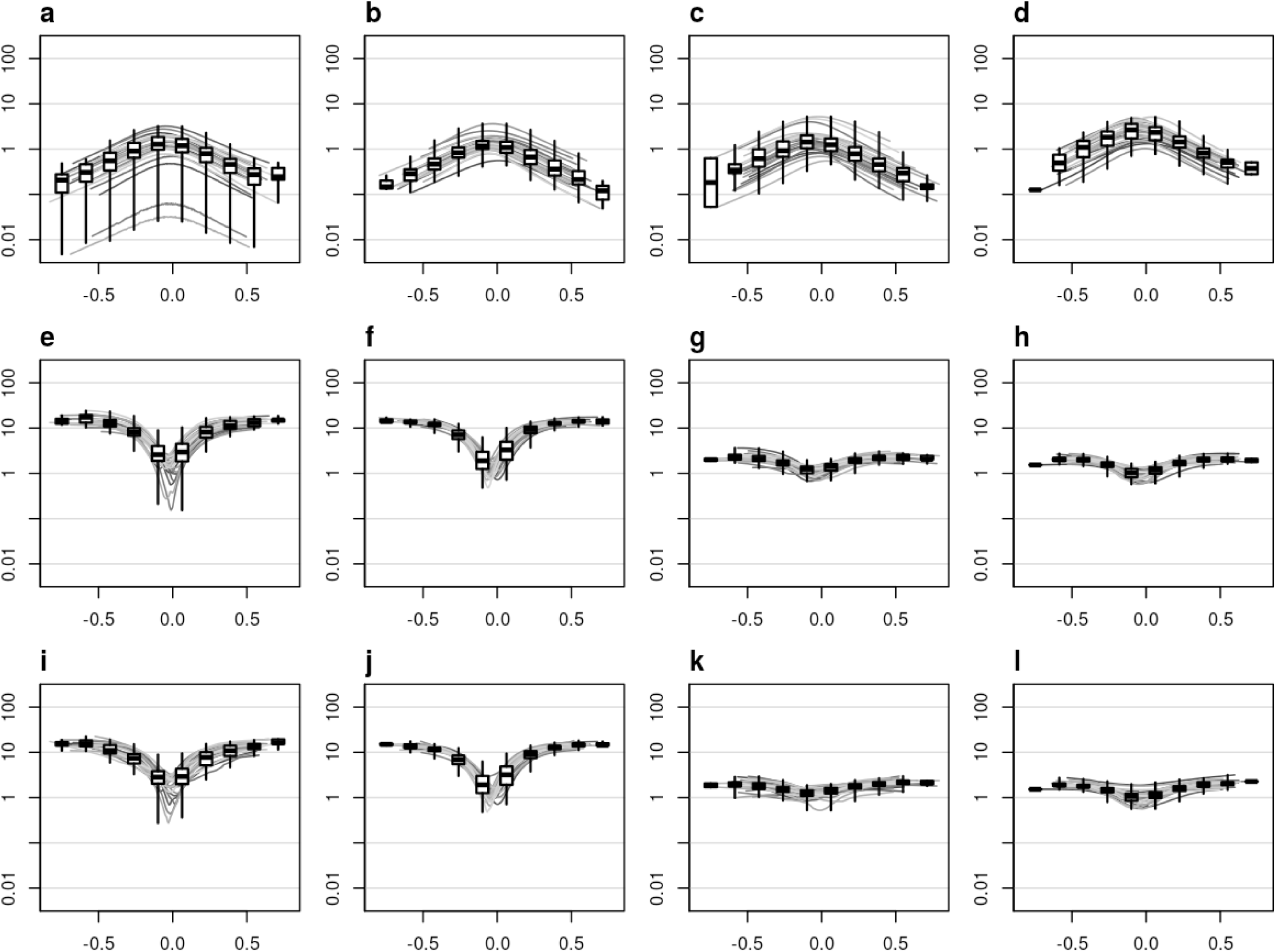
Ratio between the estimated additive variances and the true variances for each scenario: (1) *h*^2^ = 0.1 and *n* = 20 (a, e, i), (2) *h*^2^ = 0.1 and *n* = 120 (b, f, j), (3) *h*^2^ = 0.6 and *n* = 20 (c, g, k), and (4) *h*^2^ = 0.6 and *n* = 120 (d, h, l); for 100 simulations in each scenario, and for each order of random regression: order 0 (a-d), order 1 (e-h) and order 2 (i-l). Each grey curve is the ratio for 1 simulation; boxplots summarize all the point ratios in segments of a 10^th^ of the environmental gradient. The additive variances refer to the progeny population

